# Human Brain Dynamics Dissociate Early Perceptual and Late Motor-Related Stages of Affordance Processing

**DOI:** 10.1101/2023.09.07.556516

**Authors:** Sheng Wang, Zakaria Djebbara, Guilherme Sanches de Oliveira, Klaus Gramann

**Affiliations:** Biological Psychology and Neuroergonomics, Technische Universität Berlin, Berlin, Germany; Department of Architecture, Design and Media Technology, Aalborg University, Aalborg, Denmark

**Keywords:** affordances, EEG, sensorimotor time windows, time-varying automaticity, contextual dependence, physical walking & joystick movement

## Abstract

Affordances, the opportunity for action offered by the environment to an agent, are vital for meaningful behavior and exist in every interaction with the environment. There is an ongoing debate in the field about whether the perception of affordances is an automated process. Some studies suggest that affordance perception is an automated process that is independent from the visual context and bodily interaction with the environment, while others argue that it is modulated by the visual and motor context in which affordances are perceived. The present paper aims to resolve this debate by examining affordance automaticity from the perspective of sensorimotor time windows. We replicated a previous study on affordance perception in which participants actively moved through doors of different width in immersive 3D virtual environments. To investigate the impact of different forms of bodily interactions with an environment, i.e., the movement context (physical vs. joystick movement), we used the identical virtual environment from Djebbara and colleagues (2019) but displayed it on a 2D screen with participants moving through different wide doors using the keys on a standard keyboard. We compared components of the event-related potential (ERP) from the continuously recorded electroencephalogram (EEG) that were previously reported to be related to affordance perception of architectural transitions (passable and impassable doors). Comparing early sensory and later motor-related ERPs, our study replicated ERPs reflecting early affordance perception but found differences in later motor-related components. These results indicate a shift from automated perception of affordances during early sensorimotor time windows to movement context dependence of affordance perception at later stages suggesting that affordance perception is a dynamic and flexible process that changes over sensorimotor stages.

## 1 INTRODUCTION

### 1.1 Background and literature review

Affordances, the possibilities for action that the environment offers to a given organism or agent (Gibson, 1979), play a crucial role in guiding an agent’s behavior. In the field of affordance perception, there has been debate regarding whether it is an automated process or not. Some researchers argue that affordance perception is an automatic process due to its fast and effortless nature (Tucker & Ellis, 1998; Goslin, Dixon, Fischer, Cangelosi, & Ellis, 2012; Bonner & Epstein, 2017; Harel, Nador, Bonner, & Epstein, 2022), whereas others suggest that it is not automated but rather highly contextualized and can be influenced by biases and expectations (Tipper, Paul, & Hayes, 2006; Girardi, Lindemann, & Bekkering, 2010; Pellicano, Iani, Borghi, Rubichi, & Nicoletti, 2010; Kalénine, Wamain, Decroix, & Coello, 2016; Wokke, Knot, Fouad, & Ridderinkhof, 2016; Mustile, Giocondo, Caligiore, Borghi, & Kourtis, 2021). However, a synthesis perspective proposes that affordance automaticity should be understood as a dynamic process that changes over time, whereby affordance perception may initially occur automatically but is later modulated by higher-level cognitive processes (Borghi & Riggio, 2015; Kourtis, Vandemaele, & Vingerhoets, 2018; Gastelum, 2020; Djebbara et al., 2022). The question of whether affordance perception is an automated process or not may thus depend on the specific context and temporal scale being considered. The present study used brain dynamic features from electroencephalography (EEG) to provide evidence for the hypothesis that affordance perception is a time-varying process.

The question of affordance automaticity is related to the top-down/bottom-up debate regarding affordance processing (Pezzulo & Cisek, 2016) with several findings being broadly consistent with an automatic bottom up view indicating effortless processing of affordances. Using a stimulus-response compatibility (SRC) paradigm, Tucker and Ellis (1998) required participants to decide as fast as possible whether objects displayed on a computer monitor were upright or inverted. In their experiment, objects were presented in two horizontal orientations (compatible with either right-hand grasp or left-hand grasp) and two vertical orientations (upright or reverse). The authors found that participants’ responses towards upright or inverted objects were faster when the object’s handle was on the same side of the display as the response hand than when the handle was on the opposite side, although the handle of the object was irrelevant to the judgment about the objects’ vertical orientation. The results indicated that perceived affordances to grasp an object are activated automatically, even in the absence of explicit intentions to act (Tucker & Ellis, 1998). Using functional magnetic resonance imaging (fMRI), Bonner & Epstein (2017) aimed to identify representations of navigational affordances in the human visual system (Bonner & Epstein, 2017). They found that affordance information was automatically elicited during scene perception, because affordance coding in scene-selective brain regions was observed although participants performed a perceptual-semantic recognition task in which they were not explicitly asked about the navigational affordances of the scene (Bonner & Epstein, 2017). Using Bonner & Epstein’s (2017) paradigm combined with EEG, Harel et al. (2022) investigated how rapid visual information is transformed into navigationally relevant information by measuring visually-evoked event-related potentials (ERPs) with a focus on the P2 component, a positive component peaking around 220 milliseconds after stimulus onset. The authors presented participants with computer-generated room scenes featuring varying numbers of doors or paintings to manipulate the navigational affordance of the local environment. These scenes, which served as the background stimuli, were presented on the screen for 500 milliseconds. Participants were asked to report whether the horizontal or vertical bar of the central fixation cross lengthened on each trial by a factor of 25% minimizing participants’ attention to the background scenes. Although participants in the study were not required to physically move or imagine themselves moving, the authors observed that changes in navigational affordance modulated early ERPs, specifically the P2 amplitude. Harel et al. (2022) concluded that information about the potential for navigation in the scene is extracted rapidly and automatically even when participants pay no attention to the scene (Harel, Nador, Bonner, & Epstein, 2022).

Other studies have challenged the view that affordances are automatically activated, based on findings that perception of object affordances is modulated by task and context (Mazzuca et al., 2021). Context, in this case, refers to the scenario in which the object is perceived, including the presence of other objects or situations in which objects are embedded (Girardi, Lindemann, & Bekkering, 2010; Mazzuca et al., 2021; Mustile, Giocondo, Caligiore, Borghi, & Kourtis, 2021). For instance, studies by Pellicano et al. (2010) and Tipper et al. (2006) demonstrated that the affordance effects to grasp objects as proposed by Tucker and Ellis (1998) only emerge when the task requires deeper processing of the object’s characteristics, such as object categorization and shape recognition, but not in simpler perceptual tasks like color discrimination (Pellicano, Iani, Borghi, Rubichi, & Nicoletti, 2010; Tipper, Paul, & Hayes, 2006). In a behavioral study, Girardi et al. (2010) aimed to study object affordance effects in natural reach-to-grasp actions while focusing on the role of the action context in the processing of object affordances. They presented participants with a priming picture consisting of a hand and an object, and required them to make a power or precision grasp to classify the object as manmade or animate. Results showed that when the size of the object was consistent with the required grasping action (e.g., precision grasp to classify a needle as manmade), the response time was faster. Notably, the presence of this priming effect depended on the spatial arrangement of the hand and object in the picture. Specifically, the effect disappeared when the hand and object were presented as separate entities rather than an interacting pattern. Based on their findings, Girardi et al. (2010) argued that the observed affordance effect strongly depends on the action context and is not automatically activated (Girardi, Lindemann, & Bekkering, 2010). In a recent study, Mustile and colleagues (Mustile, Giocondo, Caligiore, Borghi, & Kourtis, 2021) investigated the sensorimotor brain dynamics of dangerous object perception and whether the object’s location and task goal modulated the encoding of its motor properties. Participants were recorded with high-density EEG while passively perceiving dangerous and neutral objects located at different distances from the observer and performing either a reachability judgment task or a discrimination-categorization task. The authors found that a frontal N2 potential, a component of the ERP associated with motor inhibition, was larger for dangerous objects only during the reachability judgment task, indicating task-dependent and context-dependent aversive affordances. Taken together, these studies provided evidence that affordances might not be activated automatically, because the way we look at and operate objects around us might differ depending on the meaning of that object and the context they are placed in.

Recently, a synthesis of the automated and activity-dependent views on affordance processing has emerged. According to this view, affordance perception is explained with respect to different time scales or different processing types. According to Djebbara and colleagues (2022), the sensorimotor coupling process (or sensorimotor dynamics) in enacting perception concerning affordances can be understood as changing from a macro to a micro scale, referred to as sensorimotor contingencies (SMCs; the non-specific and ongoing resonances with the environment) and sensorimotor responses (SMRs; the particular and preferential resonances with the environment occuring in the early sensorimotor processes), respectively. In line with different stages of affordance processing, Kourtis et al., (2018) found that affordances can be separated into two categories based on the temporal mechanisms of the agent-object interaction, immediate (i.e., grasping showing size-related affordances), and long-term (i.e., skillful use showing function-related affordances) with different neural representations of these two kinds of affordances (Kourtis, Vandemaele, & Vingerhoets, 2018).

### 1.2 Research question and hypothesis

The present study analyzed human brain dynamics to investigate whether affordance processing differs in automaticity over time and depending on the context. More precisely, we analyzed early and later components of evoked brain dynamics reflecting affordance processing of identical virtual spaces comparing two different movement conditions, physically walking versus joystick-controlled movements. We investigated whether different sensorimotor phases can dissociate different levels of automaticity in affordance processing dependent on the movement context. In accordance with existing research on affordance context (Girardi, Lindemann, & Bekkering, 2010; Mustile, Giocondo, Caligiore, Borghi, & Kourtis, 2021), we operationally defined context as a property of the perceived scene that primes different sensorimotor coupling states. To this end, we varied the way participants could move through a virtual environment. Varying levels of movement were realized through different displays as a between subject design, with a previous study Djebbara et al. (2019) using head mounted VR display with full body movements through the virtual environments and the present study using a 2D display with a keyboard for movement control. While moving through the identical virtual environment, the two different movement contexts should prime distinct sensorimotor coupling states. We used a virtual environment from the study by Djebbara et al. (2019) in which participants actively walked through an immersive virtual environment with the task to pass differently wide doors (passable and impassable doors) to go from one room to an adjacent room. In contrast to the physical movements through 3D virtual rooms in the Djebbara study, participants in our study used keyboard controls to move through the same rooms displayed on a 2D display. We statistically compared the ERP results from the Djebbara study with the results from the present study. Comparable early and late ERP components in both studies would support the assumption of automated affordance processing, irrespective of the movement (full-body vs. keyboard). Differences in both, early and late ERP components, in contrast, would support the assumption of context dependent affordance processing while comparable early but differences in late components would support a stage-dependent processing of affordances.

## 2 METHODS

### 2.1 Participants

Eighteen participants (7 females) without history of neurological pathologies were recruited from social media and a participant pool of the Technical University of Berlin, Germany. All participants signed the consent form, which was approved by the Ethics Committee of the Technical University of Berlin. After each data collection, participants received either monetary compensation (10€/hour) or accredited course hours. Their mean age was 24.0 years (σ = 3.8). All participants were right-handed, had normal or corrected-to-normal vision, and none had a specific background in architecture (i.e., no architects or architectural students). We excluded one male participant from the study because of experienced dizziness.

### 2.2 Experimental Paradigm and Procedure

We used the same experimental paradigm and stimuli as Djebbara et al., (2019, 2021) and repeated their experiment in a stationary desktop setup instead of physically walking through a virtual space with head mounted VR. The experiment was conducted in a dimly lit experimental room at the Berlin Mobile Brain/Body Imaging Laboratories (BeMoBIL) using a 27’’, LED monitor (PROLITE XB2780HSU-B1 (EOL), full HD 1920 × 1080 resolution, horizontal sync 24 - 80kHz, and vertical sync 56 - 70Hz) to display the virtual environment. The size of the virtual space was 9 m × 5 m (virtual meters), with room sizes of 4.5 m × 5 m for two adjacent rooms connected through a door (see Fig. 1). Participants were seated in front of the monitor in a comfortable position and their task was to move through the virtual environment using four arrow keys on the keyboard. The doors connecting the two rooms differed in width from wide (1.5 m, easily pPassable) to mid (1 m, passable) to narrow (0.2 m, unpassable) in a pseudorandomized order. The door width varied to manipulate the transition affordance between rooms. Participants performed a forewarned (S1-S2) Go/NoGo paradigm (pseudorandomized 50/50) that required them to walk from one room to a second room and to pick up a token in half of the trials while in the other half of the trials they simply judged the emotional valence of the first room they were located in without moving to the second room.

**Fig. 1.**
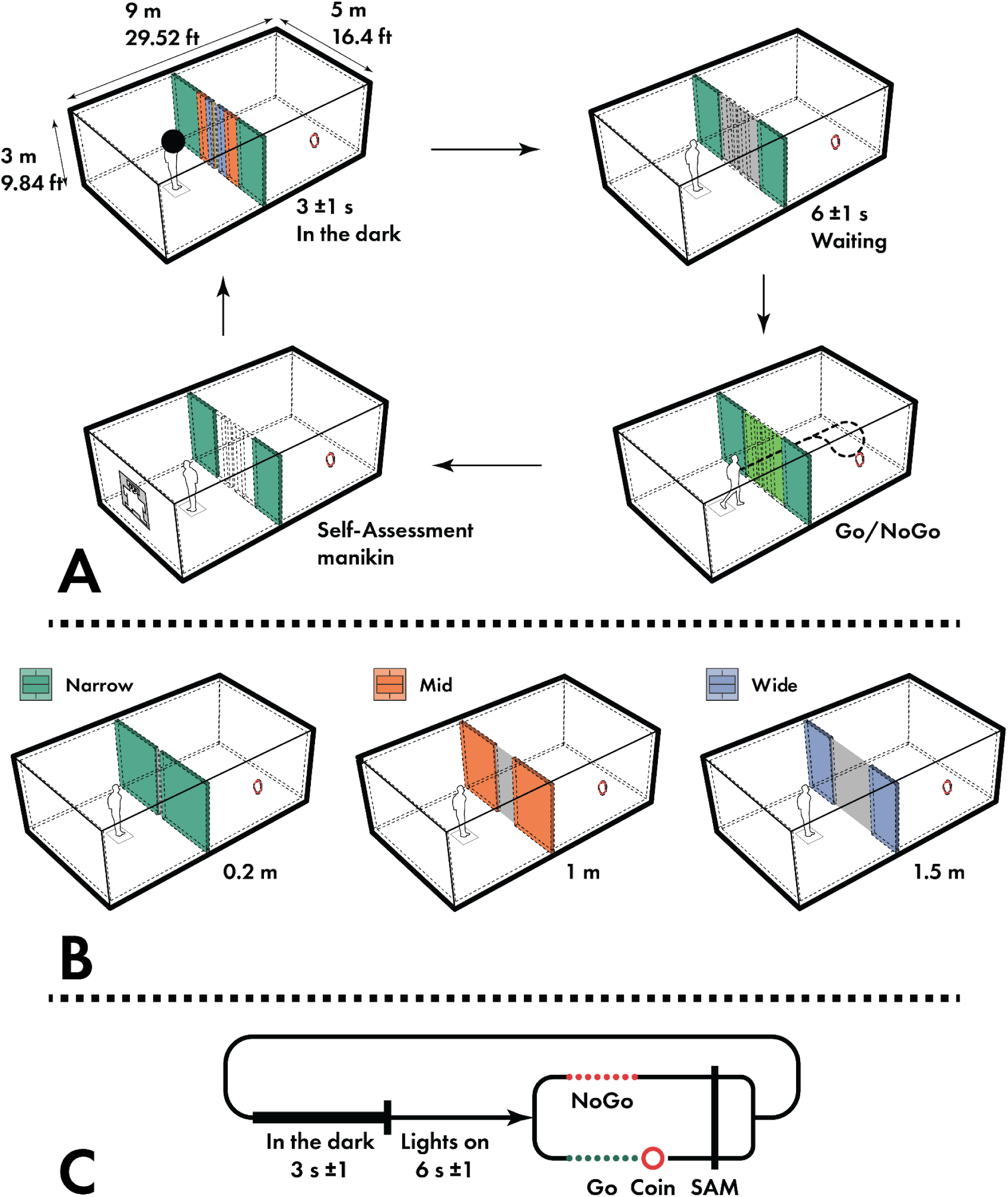
(*A*) Participants were instructed to view the screen. First, in the start square within the Unity environment, the screen was pure black for 3 s (σ = 1 s), because of a black sphere restriction within Unity. Second, the moment the light was on and the black sphere disappeared, participants perceived the door they had to pass. The door width was either narrow (0.2 m), mid (1 m) or wide (1.5 m). Then they waited for the imperative stimulus, either a green door (Go) or a red door (NoGo), for 6 s (σ = 1s). In the case of Go, participants were instructed to walk toward the door, using the keyboard keys. Upon entering the second room, participants had to find and virtually touch a red rotating coin. Successfully touching the coin earned participants an additional 1 cent as a bonus. The bonus number was displayed in the top corner of the screen and updated regularly. Participants then returned to complete the virtual SAM questionnaire. In the case of NoGo, participants were instructed to turn around and complete the virtual SAM questionnaire. Finally, after completing the SAM questionnaire, participants were instructed to press the SPACE key to restart the next trial. (*B*) There were 3 dimensions of door width conditions: Narrow, 0.2 m (green); Mid, 1 m (red); Wide, 1.5 m (purple). Note the color code for each door as used throughout the paper. (*C*) A diagrammatic timeline depicting the sequence of events for a single trial.

The experiment was thus a 2 × 3 repeated-measures design including the factors imperative stimulus (Go, NoGo) and door width (Narrow, Mid, Wide). For each participant, there were 240 trials in total, with 40 trials for each condition. In each trial, participants started in a dark environment on a predefined starting square (Fig. 1). After a random intertrial interval (mean = 3 s, SD = 1 s), the “lights” were turned on, and participants viewed a room with a closed door. After waiting for a random duration (mean = 6 s, σ = 1 s), the door color changed into either green indicating a Go trial or red indicating a NoGo trial. If the door color was green, participants were instructed to move toward the door by pressing the arrow keys. Once they reached the door, the door slid open aside immediately. Participants were instructed to find and virtually touch a red rotating ring in the second room. Touching the ring earned participants 1 cent as a bonus to the hourly rate, which was also our incentive strategy. After participants went back to the first room, they were instructed to return to the starting square and to provide an emotional rating using the Self-Assessment Manikin (SAM) questionnaire. However, in the case of a red door, participants did not walk through the transition but directly provided an emotional rating for the room they were located in. After the emotional rating, participants pressed the SPACE bar to start the next trial.

In each Go trial, participants were instructed to move through the door into the second room irrespective of whether the door was passable or not passable. This setting kept motor processes comparable for all Go trials irrespective of the transition affordance (passable or not passable). When participants touched the surrounding walls, the walls turned red, indicating that they failed to pass and the red rotating ring located at the second room would disappear indicating that participants did not get a bonus for this trial. Participants first finished a training session to become accustomed to the experimental environment and the different conditions. The experimenter communicated with participants from a control room, separated from the experimental space, using a camera and a microphone-speaker system. Moreover, there were 5 breaks in total, including 2 mandatory breaks (2 minutes at least) and 3 optional breaks.

In order to investigate the potential impact of movement on affordance perception, we compared the ERP results with the findings of Djebbara et al. (2019). This allowed for a direct comparison of the same visually presented transitional affordances in two different movements (physical walking vs. keyboard movement). In addition, we administered the Igroup Presence Questionnaire (IPQ) before and after the experiment to assess participants’ sense of presence (Slater & Sanchez-Vives, 2016). The IPQ consists of 4 dimensions: general presence (PRES), spatial presence (SP), involvement (INV) and experienced realism (REAL) (Grassini & Laumann, 2020; Schubert, Friedmann, & Regenbrecht, 2001). The data were analyzed using paired-samples t tests with the factor time with two levels (pre- and post-test) as a within-subjects factor.

### 2.3 Self Assessment Manikin

To investigate the subjective experience of the task, we asked participants to finish a virtual SAM questionnaire after each trial. The SAM is a pictorial assessment of pleasure, arousal, and dominance on a 5-point Likert scale (Bradley & Lang, 1994). Participants were asked to self-assess their current state after each trial by pressing the respective keyboard number on the 5-point scale. A first key press selected participants’ answers but still allowed changes to the selection. By pressing the same key again, the response was confirmed. The SAM data were analyzed using a 2 × 3 factorial repeated-measures ANOVA with the factors imperative stimulus (Go and NoGo), door width (Narrow, Mid, and Wide). For post hoc analysis, the data were contrasted using least significant difference (LSD) confidence interval adjustment.

### 2.4 EEG Recording and Data Analysis

Experimental stimuli based on Unity software were displayed on a standard monitor. All data streams, including experimental events such as participants’ movement and location in the Unity environment, stimulus markers and key presses, SAM answers and EEG data, were recorded and synchronized using LabStreamingLayer (LSL; Kothe, Medine, Boulay, Grivich, & Stenner, 2014).

EEG data were recorded with a 64-channel EEG system (ANT Neuro), sampled at 500 Hz and referenced to CPz. Impedances were kept below 10 kΩ. Data analysis was conducted using MATLAB (MathWorks), the EEGLAB toolbox (Delorme & Makeig, 2004), and custom scripts. Before data preprocessing, all training and resting data sessions were excluded. All preprocessing steps were based on the BeMoBIL pipeline (Klug et al., 2022), first down-sampling the data to 250 Hz and then applying the ZapLine-plus function to remove spectral peaks at 50 Hz, corresponding to the power line frequency (Klug and Kloosterman, 2022). Next, bad channels were detected and interpolated in a repeated process using the clean raw data plugin of EEGLAB. To ensure a reproducible bad channel detection, we ran the *clean_artifacts function*, with a recommended minimum of 10 iterations. For the automatic channel cleaning, the data was split into time windows of five seconds and robust interpolations of each channel were computed. Channels with a correlation to a random sample of other channels were marked for removal in case the correlation was below 0.8 for more than 50% of the processed data and subsequently interpolated using a spherical spline interpolation. As a final step, the data was re-referenced to the average of all scalp channels, excluding the EOG channel. Subsequently, on each cleaned dataset, we computed an independent component analysis (ICA) using an adaptive mixture independent component analysis (AMICA) algorithm (Palmer, Kreutz-Delgado, & Makeig, 2012) with the recommended parameter values from Klug and Gramann (2021). Before computing ICA, we performed high-pass filtering with a 0.75 Hz cutoff frequency. Then the AMICA decomposition was computed using one model and 10 iterations of data rejection with a rejection threshold of 2.9 standard deviations from the log likelihood of the explained data during the decomposition. Afterwards, an equivalent current dipole model was computed using the *DIPFIT* toolbox of EEGLAB. Independent components (ICs) were automatically classified using ICLabel (Pion-Tonachini, Kreutz-Delgado, & Makeig, 2019). Subsequently, we removed ICs reflecting eye-movement activity. The computed AMICA information including rejections and dipole fitting was copied back to the initial preprocessed, but unfiltered, dataset.

The continuous datasets were bandpass-filtered between 0.2 Hz and 40 Hz and two sets of epochs were extracted with the first set for the onset of the first room and the second set with onset of the imperative stimulus. All epochs were extracted from −1000 ms before to 1000 ms after stimulus onset for Narrow, Mid, and Wide door trials separately. Then, all epochs were baseline corrected using a -200ms and 0 ms pre-stimulus interval and cleaned using the *bemobil_reject_epochs function* to rank epochs with respect to their noise level and removing a fixed percentage of the worst ranking epochs (Klug et al., 2022). Each epoch was evaluated using four measures that were normalized by their median across epochs: i) mean of channel means (weight = 1), ii) mean of channel SDs (weight = 1), iii) SDs of channel means (weight = 1), and iv) the SD of channel SDs (weight = 1). Each epoch then received a final summed score and the epochs were sorted according to that score. Finally, up to 10% of epochs were removed (mean = 23.62, SD = 0.697) accordingly.

Early perceptual components of the ERPs time-locked to the onset of the virtual room (lightsOn) were analyzed at midline posterior leads (Pz, POz, and Oz) comparing ERPs from Djebbara and colleagues (2019) with the current results. ERPs time-locked to the subsequent onset of the imperative stimulus (Go/NoGo) were analyzed at the same leads with the inclusion of one additional anterior electrodes (Fz, FCz, and Cz) to further allow for a direct statistical comparison of the motor-related ERPs of Djebbara et al. (2019) and the corresponding data of the current study. The results of the anterior leads focus on FCz as described in Djebbara et al. (2019)(additional results for the other two anterior electrodes Fz and Cz are provided in the supplement). These electrodes locations cover brain regions involved in processing visual information and regulating motor actions as used in previous studies (Bozzacchi, Giusti, Pitzalis, Spinelli, & Di Russo, 2012; Bozzacchi, Spinelli, Pitzalis, Giusti, & Di Russo, 2015; Djebbara et al., 2019; Mustile, Giocondo, Caligiore, Borghi, & Kourtis, 2021; Harel, Nador, Bonner, & Epstein, 2022). We identified three peaks (N120, P164, and N260) after the “lights on” stimulus onset through visual inspection at the grand average level. Individual peak latencies were determined considering both EEG polarity and a time range of -50 ms to +50 ms around the group-level peak latencies. Therefore, the search windows for individual peaks ranged from 70 to 170 ms, 114 to 214 ms, and 210 to 310 ms for the N120, P164, and N260 peaks, respectively. To ensure robust statistical analysis, each negative or positive peak was exported, computing the mean from the data covering the peak plus/minus 3 sample points before and after the peak. The data of P164 and N260 were analyzed using a 2 × 3 mixed-measures ANOVA with a between-subjects factor movement (physical walking vs. keyboard movement), and a within-subjects factor door width (Narrow, Mid, and Wide). For post hoc analysis, the data were contrasted using Tukey’s HSD and LSD confidence interval adjustment. Because the N120 component was an additional negative peak detected in the present study that was absent in the study of Djebbara et al. (2019), we conducted an additional one-way ANOVA with the door width (Narrow, Mid, and Wide) as a repeated-measures factor and the amplitude of the N120 as dependent measure. For post hoc analysis, the data were contrasted using LSD confidence interval adjustment.

For the later motor-related cortical potentials (MRCPs), we focused on the early postimperative component (EPIC) and postimperative negative variation (PINV) as reported in Djebbara and colleagues (2019). After the stimulus onset of the imperative (Go/NoGo) stimulus, a negative peak at posterior leads was observed around 236 ms and a positive peak at FCz electrode was observed around 280 ms, referred to as the early postimperative component (EPIC) (Djebbara et al., 2019). Search windows for individual peaks were ranging from 186 ms to 286 ms and 230 ms to 330 ms respectively. For the sake of robust statistics, peak amplitudes were calculated including the peak plus/minus 3 sample points before and after this detected peak. The PINV can be considered a postimperative negative variation (PINV) reflecting the immediate motor execution related to onset of an imperative stimulus (Casement et al., 2008; Djebbara et al., 2019; Elbert, Rockstroh, Lutzenberger, & Birbaumer, 1982). As in the previous study of Djebbara et al. (2019), the PINV component was calculated as the mean amplitude in the range of 600-800 ms after stimulus onset (showImperative: Go/NoGo). The data of EPIC and PINV were both analyzed using a 2 × 2 × 3 mixed-measures ANOVA with a between-subjects factor movement (physical walking and keyboard movement), two within-subjects factors imperative stimulus (Go and NoGo), and door width (Narrow, Mid, and Wide). For post hoc analysis, the data were contrasted using Tukey’s honest significant differences (HSD) and LSD confidence interval adjustment.

## 3 RESULTS

In the current study, we investigated whether affordance perception of architectural transitions (passable and impassable doors) is similar for full-body walking as compared to keyboard-controlled movements through the identical environment to provide further insights into whether affordance processing is automated or not. To this end, we analyzed and statistically compared electrocortical activity from two studies in which participants moved through the identical virtual environment either physically or based on keyboard controls.

### 3.1 IPQ Data

In this study, the IPQ questionnaire was used before and after the main experiment in order to investigate the extent of participants’ sense of presence. A paired sample t test for the pre- and post-test data for each dimension of the IPQ revealed significant differences in general presence (PRES) (T_16_ = -5.191, P < 0.001, Cohen’s D = -1.259), spatial presence (SP) (T_16_ = -6.014, P < 0.001, Cohen’s D = -1.459), involvement (INV) (T_16_ = -2.507, P = 0.023, Cohen’s D = -0.608), and experienced realism (REAL) (T_16_ = -3.228, P = 0.005, Cohen’s D = -0.783) (see Fig. 2).

**Fig. 2.**
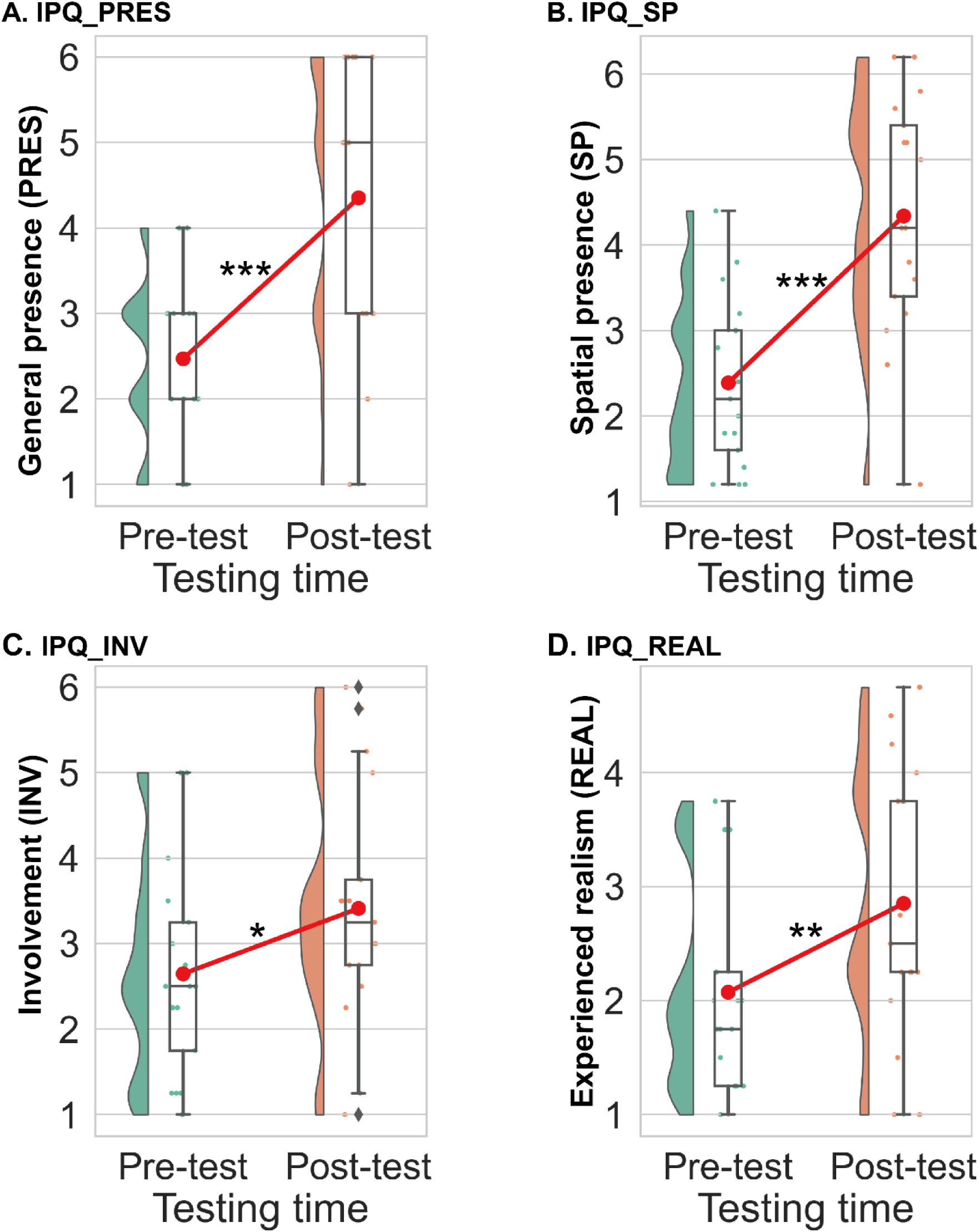
Raincloud plots displaying the distribution of IPQ ranking scores for both the pre-testing and post-testing conditions. Paired-sample t-tests were conducted to analyze the four dimensions of the IPQ questionnaire. The means for each dimension are indicated by red points and solid lines. The medians are indicated by black solid lines on each boxplot. Statistical significance was denoted as *P < 0.05, **P < 0.01, and ***P < 0.001.

### 3.2 Subjective Data: SAM Ratings

The SAM questionnaire was completed for both Go and NoGo trials in all door width conditions. A 2 × 3 factorial repeated-measures ANOVA was used with the factor imperative stimulus (Go and NoGo) and the factor door width (Narrow, Mid, and Wide) for each emotional dimension of the SAM questionnaire (see Fig. 3). For Arousal, there was a significant main effect of the factor door width (F_2,32_ = 9.628, P < 0.001, partial η^2^ = 0.376) as well as a significant main effect of the factor imperative stimulus (F_1,16_ = 53.061, P < 0.001, partial η^2^ = 0.768). The interaction of both factors was significant (F_2,32_ = 19.106, P < 0.001, partial η^2^ = 0.544). Post-hoc comparisons revealed that only the arousal score of medium wide doors was significantly different from the arousal score of wide doors (P = 0.043) in the NoGo condition. However, in the Go condition, significant differences were identified for the comparison of Narrow vs. Wide doors (P < 0.001), Narrow vs. Mid doors (P < 0.001), and Mid vs. Wide doors (P = 0.021). For Dominance, there was a significant main effect of the factor door width (F_2,32_ = 13.944, P < 0.001, partial η^2^ = 0.466) as well as a significant main effect of the factor imperative stimulus (F_1,16_ = 21.198, P < 0.001, partial η^2^ = 0.570). Both factors interacted significantly (F_2,32_ = 19.878, P < 0.001, partial η^2^ = 0.554). Post hoc comparisons revealed no differences in dominance ratings between differently wide doors in the NoGo condition. However, in the Go condition, significantly lower ratings were observed for the Narrow as compared to Wide doors (P < 0.001), Narrow as compared to Mid doors (P < 0.001), and a tendency towards a difference for the comparison of Mid vs. Wide doors (P = 0.061). These results were also found for the Valence dimension of the SAM including significant main effects of the factor door width (F_2,32_ = 33.737, P < 0.001, partial η^2^ = 0.678) and imperative stimulus (F_1,16_ = 46.657, P < 0.001, partial η^2^ = 0.745) as well as their interaction (F_2,32_ = 48.585, P < 0.001, partial η^2^ = 0.752). The three door widths levels did not differ significantly under the NoGo condition. However, under the Go condition, significant differences were identified in Narrow vs. Wide (P < 0.001), Narrow vs. Mid (P < 0.001) and Mid vs. Wide doors (P = 0.027).

**Fig. 3.**
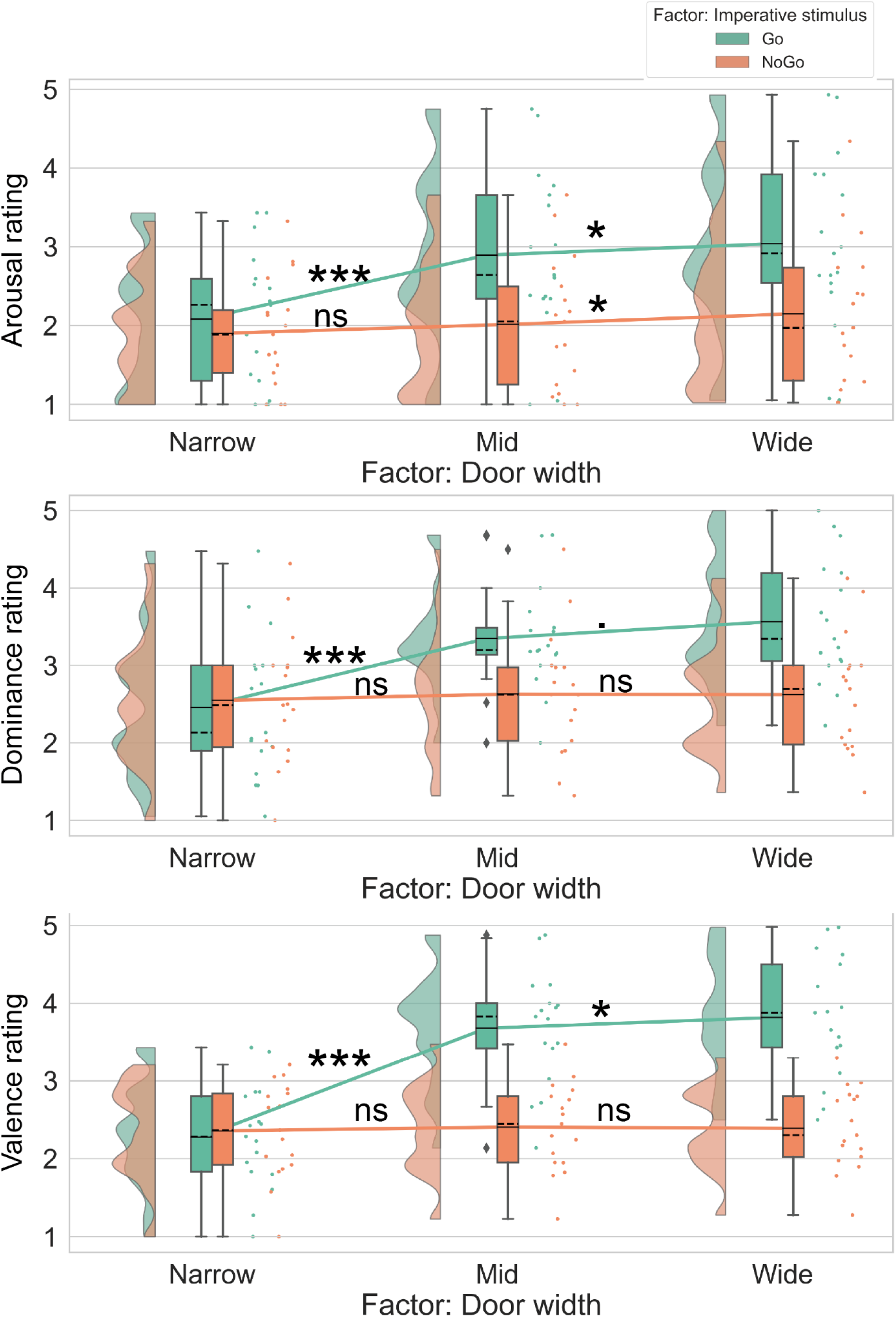
Raincloud plots displaying SAM rating for two factors: door width (Narrow, Mid and Wide) and imperative stimulus (Go, NoGo). The means are indicated by solid lines, medians, by dashed lines. Statistical significance was denoted by ▪P < 0.1, *P < 0.05 and ***P < 0.001, while non-significant results were denoted by ’ns’.

### 3.3 EEG data: Early ERPs

We hypothesized that ERP components in both studies should be comparable during early sensorimotor stages if affordance processing was an automated process, irrespective of the movement condition. To test this assumption, we analyzed the early ERP components reflecting perceptual aspects of transition affordances from the present study and compared the results with the results reported by Djebbara et al., (2019). For ERPs with onset of the rooms including different door widths, three prominent peaks were observed over posterior leads. Over the occipital midline electrode, a first negative component peaked around 120 ms, followed by a positive peak around 164 ms and a negative peak around 260 ms (Fig. 4). The first negative component was not observed in the study by Djebbara et al. (2019) and thus, no comparison between the movement conditions was possible. A statistical test for this N120 component was conducted only for the desktop condition while the comparison of visual evoked potentials from Djebbara et al. and the present study was computed for the latter two components using a mixed-measures ANOVA with the between-subjects factor movement (physical walking vs. keyboard movement) and the within-subjects factor door width (Narrow, Mid, and Wide) (Fig. 5).

**Fig. 4.**
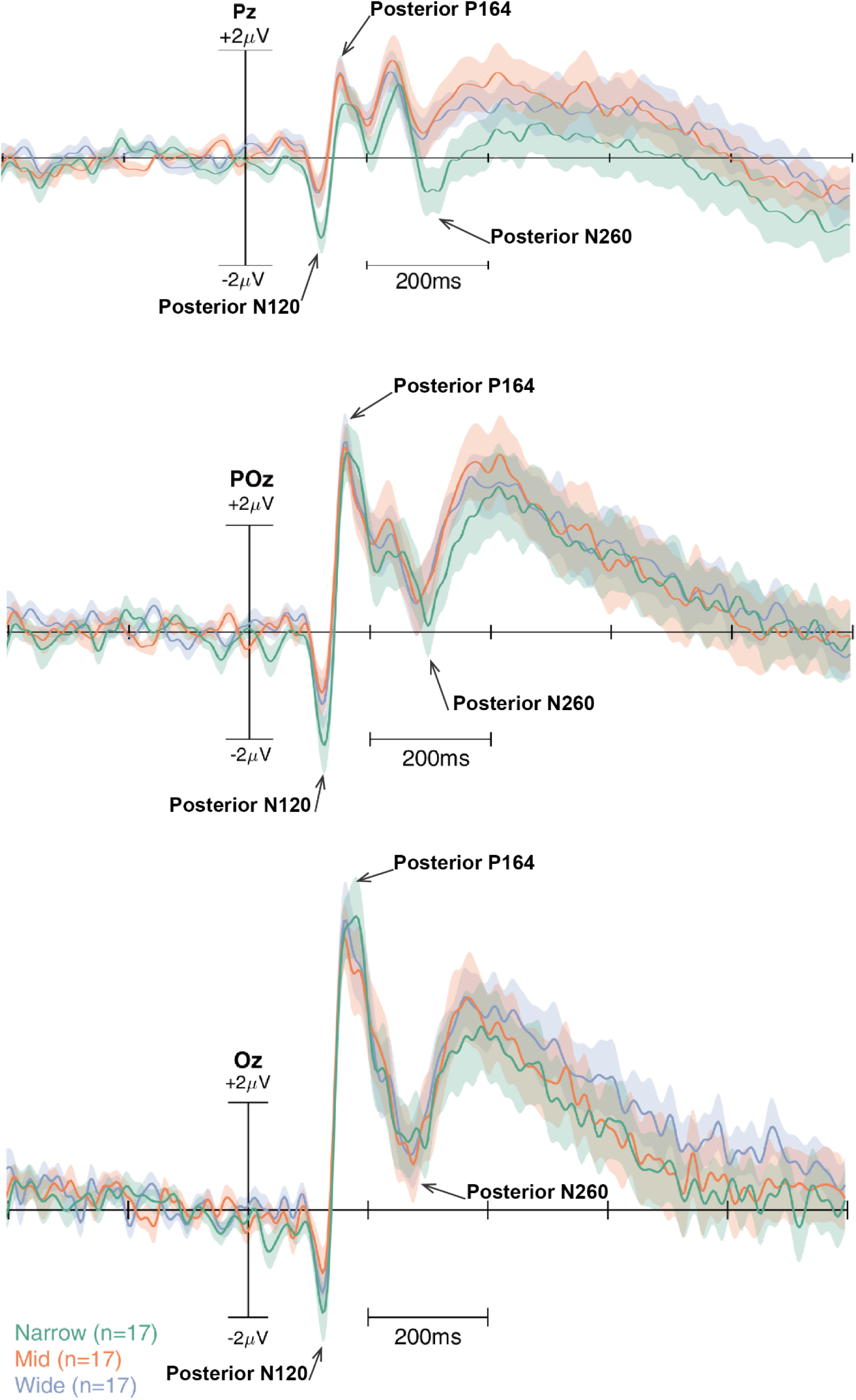
Grand average event-related potentials with onset of the room display at posterior leads Pz, POz and Oz in the keyboard movement condition. The different door widths are color coded with the Narrow door condition in green, the Mid door condition in red, and the Wide door condition in purple. Three prominent components (N120, P164, N260) as observed over electrodes Pz, POz and Oz are indicated by arrows.

**Fig. 5.**
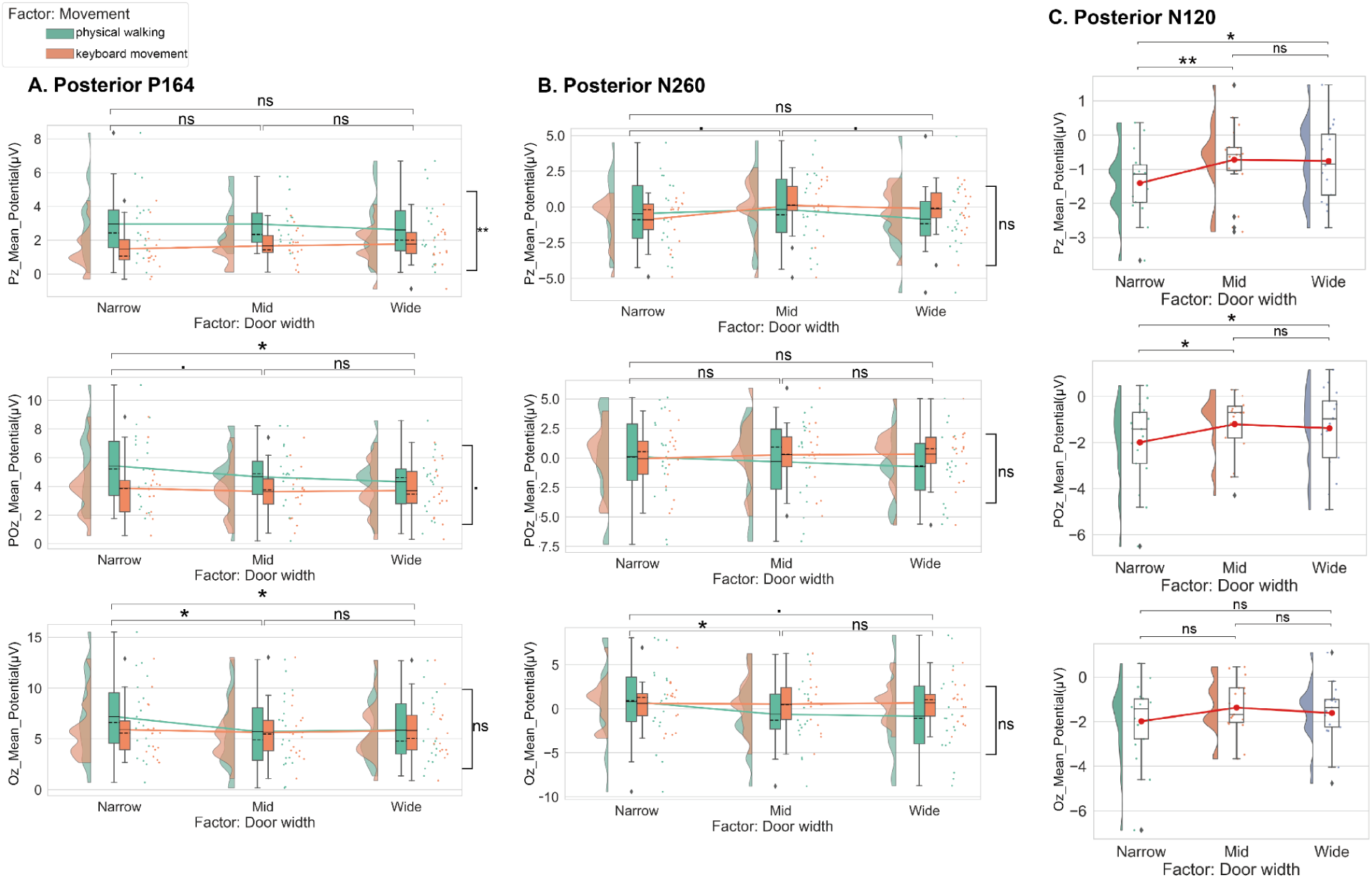
(A & B) Raincloud plots displaying mean amplitudes of the two early ERP peaks (P164, N260) at posterior electrodes for three different door width conditions (Narrow, Mid, and Wide) and two movement conditions; (C) and also mean amplitudes of the early ERP peak (N120) at posterior electrodes for three different door width conditions, detected at the present study additionally. For raincloud plots A & B, the means are indicated by solid lines; medians, by dashed lines. For raincloud plots C, the means are indicated by red solid points; medians, by black solid lines. Results from multiple comparisons using the Tukey’s HSD test and LSD adjustment are indicated with stars. Statistical significance was denoted by ▪P < 0.1, *P < 0.05, **P < 0.01, and ***P < 0.001, while non-significant results were denoted by ’ns’.

#### Early posterior positive component around 164 ms

The P164 component observed in the present study showed similar topographies and was closest in latency to the P140 of Djebbara et al. (2019). For electrode Pz, we found no influence of the door width on P164 amplitudes (F_2,68_ = 0.110, P = 0.896), but a main effect of the factor movement (F_1,34_ = 8.000, P = 0.008, partial η^2^ = 0.190) and no interaction between the two factors (F_2,68_ = 0.825, P = 0.443). For the parieto-occiital lead (POz), we found a main effect of the factor door width (F_2,68_ = 3.355, P = 0.041, partial η^2^ = 0.090), and a slight tendency of the main effect of the movement condition (F_1,34_ = 2.900, P = 0.098) but no interaction of the two factors (F_2,68_ = 1.672, P = 0.196). Post hoc comparisons revealed no differences in P164 amplitudes between Mid and Wide doors (P = 0.496), but a trend between Narrow and Mid doors (P = 0.092) and significant difference between Narrow and Wide doors (P = 0.028). Finally, for Oz, there was a main effect of the factor door width on the amplitude of the P164 component (F_2,68_ = 4.060, P = 0.022, partial η^2^ = 0.107), but no main effect of the factor movement (F_1,34_ = 0.215, P = 0.646), and a slight tendency of the interaction of both factors (F_2,68_ = 2.389, P = 0.099). Post hoc comparisons for the different door width conditions showed no amplitude differences between Mid and Wide doors (P = 0.567), but significant differences between Narrow and Mid (P = 0.022), and between Narrow and Wide (P = 0.027) doors.

#### Early posterior negative component around 260 ms

Because of similarities in the topography and latency of the observed N260 in the present study with the N215 from Djebbara et al. (2019), we compared the two ERP components statistically. For Pz, we found a marginally significant main effect of the factor door width (F_2,68_ = 2.569, P = 0.084, partial η^2^ = 0.070). There was no main effect of the factor movement (F_1,34_ = 0.094, P = 0.761) and no interaction effect between the two factors (F_2,68_ = 1.856, P = 0.164). For POz, there was no main effect of the factor door width (F_2,68_ = 0.355, P = 0.702), no main effect of the factor movement condition (F_1,34_ = 0.286, P = 0.596) or an interaction between the two factors (F_2,68_ = 2.060, P = 0.135). Over the occipital lead (Oz), we found a marginally significant main effect of the factor door width (F_2,68_ = 2.539, P = 0.086, partial η^2^ = 0.069), no main effect of the factor movement (F_1,34_ = 0.522, P = 0.475) and a tendency of an interaction effect of both factors (F_2,68_ = 2.646, P = 0.078).

#### Desktop Condition Component Posterior N120

The N120 was an additional negative component observed in our keyboard movement condition. Therefore, the N120 amplitude of each posterior electrode was calculated using one-way ANOVA with repeated measures for door widths. For Pz, differences of N120 amplitudes on three door widths were found (F_2,32_ = 4.300, P = 0.022, partial η^2^ = 0.212). Post hoc multiple comparisons showed that significant differences were identified for the comparison of Narrow vs. Wide doors (P = 0.025), Narrow vs. Mid doors (P = 0.008), but no significant differences were found for the comparison of Mid vs. Wide doors (P = 0.900). For POz, differences of N120 amplitudes on three door widths were found (F_2,32_ = 4.391, P = 0.021, partial η^2^ = 0.215). Post hoc comparisons showed no significant differences for the comparisons of Mid vs. Wide doors (P = 0.525), but significant differences for the comparison of Narrow vs. Wide doors (P = 0.045) and Narrow vs. Mid doors (P = 0.015). For Oz, there were no significant differences of N120 amplitudes on three door widths (F_2,32_ =1.790, P = 0.183).

### 3.4 Motor-related ERPs

Besides assuming comparable early ERPs to reflect automatic affordance perception, we further hypothesized that in the subsequent motor-related stage of sensorimotor coupling, the movement context might moderate late ERP components reflecting later movement context-dependent affordances. In order to test this assumption, we directly compared the motor-related potentials of transition affordances in the present study with the corresponding results of Djebbara et al., (2019). The early postimperative component (EPIC) and postimperative negative variation (PINV) were observed at central midline leads (FCz, Pz, POz, and Oz) after the onset of the imperative stimulus, which instructed participants to go or not to go through the door to the next room (Fig. 6). Both the posterior EPIC components and anterior EPIC (at electrode FCz) are displayed in Figure 7.

**Fig. 6.**
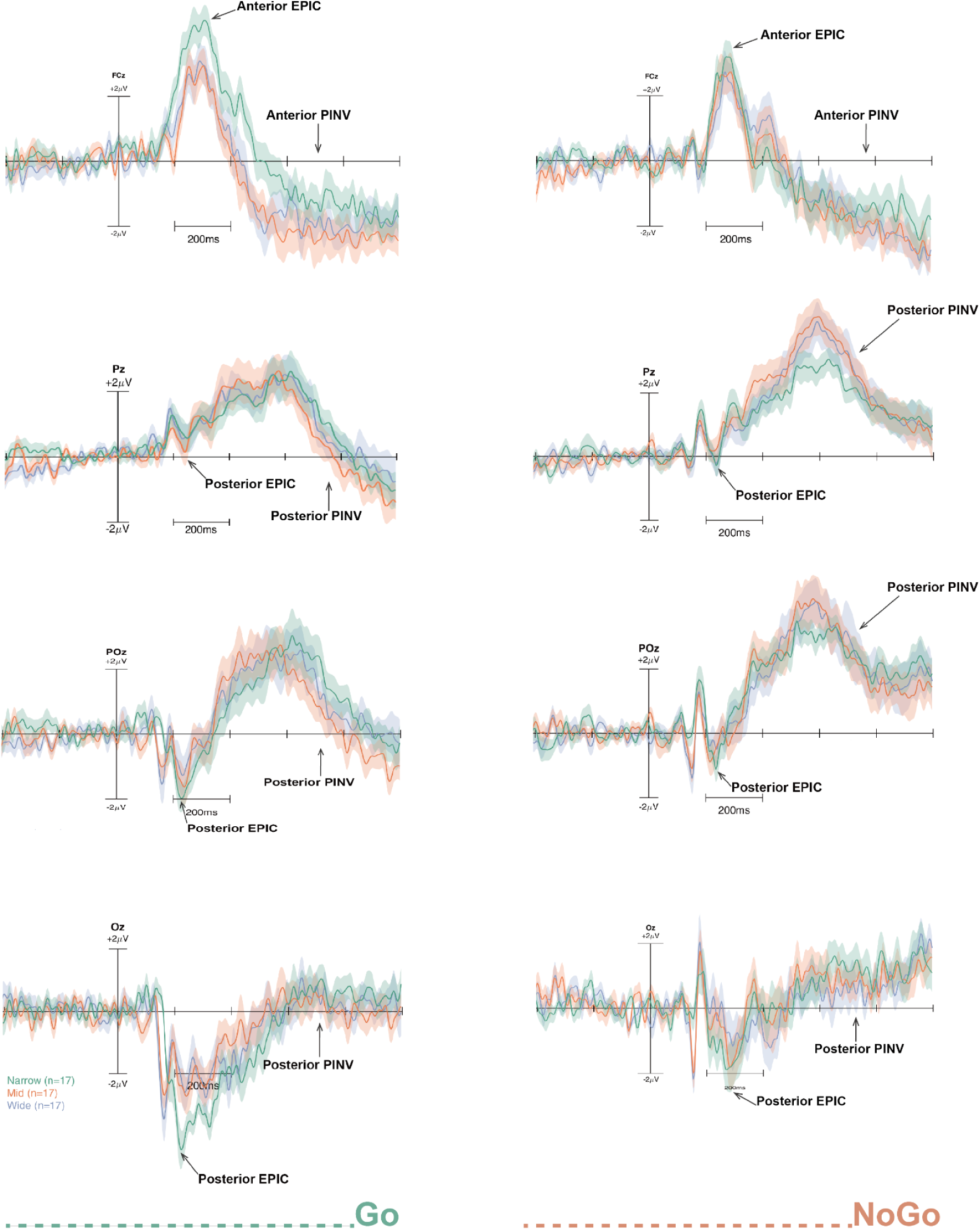
Six time-locked event-related potentials (ERPs) from electrode FCz, Pz, POz, and Oz at the onset of imperative stimulus (Go, NoGo) under the present keyboard movement condition. The Narrow condition is in green, the Mid condition is in red, and the Wide condition is in purple. The plots on the left column showed the ERPs under the Go condition, while the plots on the right column showed the ERPs under the NoGo condition. Two components (EPIC, and PINV) were marked with arrows.

**Fig. 7.**
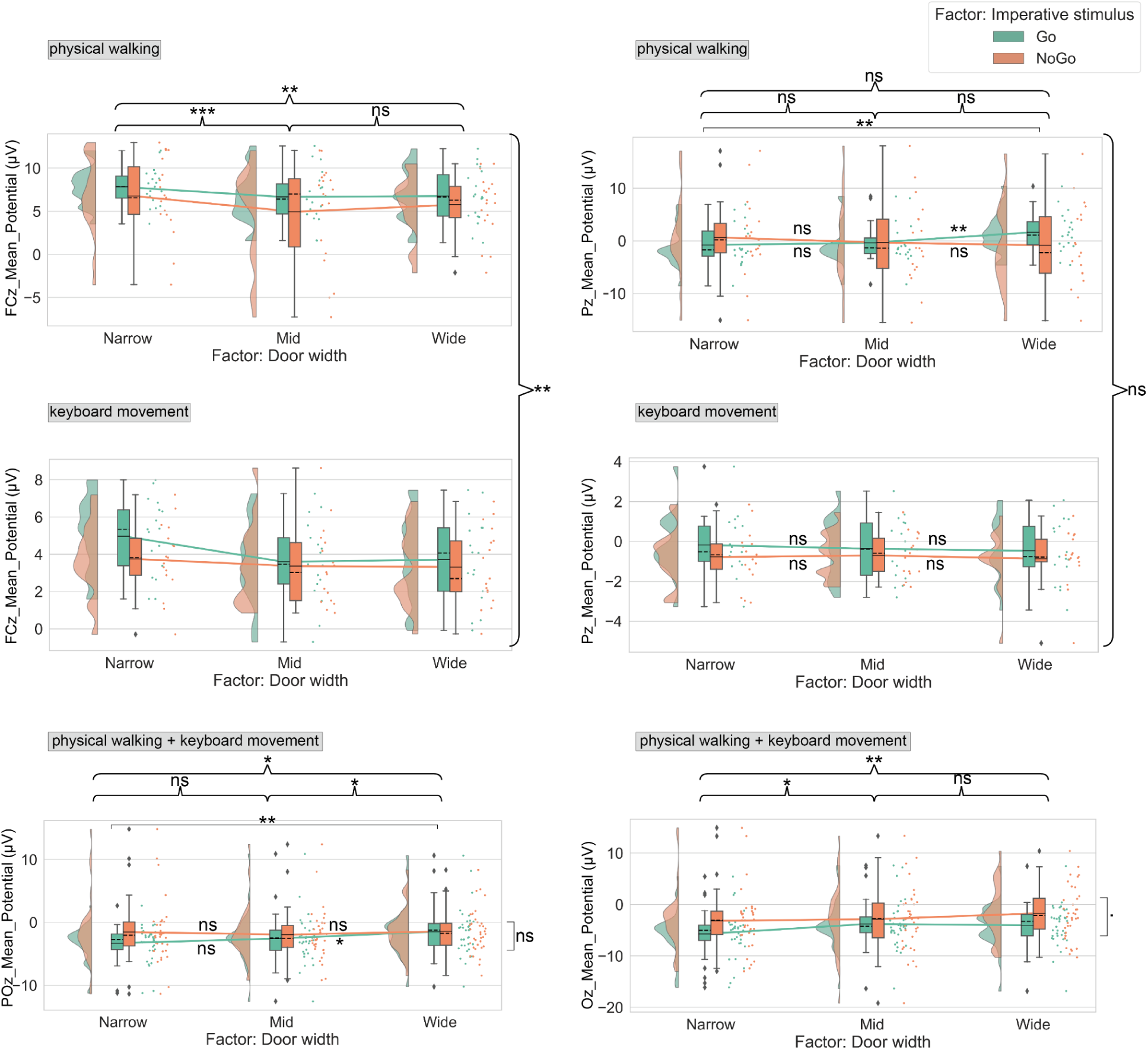
Raincloud plots displaying mean amplitudes of EPIC peaks at anterior and posterior electrodes (FCz, Pz, POz, and Oz) for three factors (three different door width conditions (Narrow, Mid, and Wide), imperative stimulus (Go, NoGo) conditions and two movement conditions). The means are indicated by solid lines; medians, by dashed lines. Results of the main effect of the factor door width are indicated by braces, while results of the interaction effect are indicated by square brackets or directly on the solid lines. Results from multiple comparisons using the Tukey’s HSD test and LSD adjustment are indicated with stars. Statistical significance was denoted by ▪P < 0.1, *P < 0.05, **P < 0.01, and ***P < 0.001, while non-significant results were denoted by ’ns’.

#### 3.4.1 EPIC

##### Anterior positive EPIC - Comparing physical walking and keyboard movement

For the anterior EPIC, we conducted a mixed-measures ANOVA with a between-subject factor movement (physical walking vs. keyboard movement) and two within-subject factors with 3 (door width) and 2 (imperative stimulus) levels for the electrode FCz. We found a main effect of the factor door width (F_2,68_ = 8.238, P < 0.001, partial η^2^ = 0.195), a tendency for a main effect of the factor imperative stimulus (F_1,34_ = 3.481, P = 0.071, partial η^2^ = 0.093) and a main effect of the factor movement (F_1,34_ = 11.178, P = 0.002, partial η^2^ = 0.247). The mean amplitudes of the EPIC in the physical walking condition were significantly larger than those in the keyboard movement condition. None of the relevant interactions reached significance (imperative stimulus × door width: F_2,68_ = 0.329, P = 0.721; three-way interaction: F_2,68_ = 1.103, P = 0.338). Post hoc comparisons for the different door width conditions showed significant differences for the comparison of Narrow vs. Mid doors (P < 0.001), and Narrow vs. Wide doors (P = 0.005), but not for the comparison of Mid vs. Wide doors (P = 0.429).

##### Posterior negative EPIC - Comparing physical walking and keyboard movement

For the posterior EPIC, we conducted a mixed-measures ANOVA with the between-subject factor movement context (physical walking vs. keyboard movement) and two within-subject factors with 3 (door width) and 2 (imperative stimulus) levels separately for electrodes Pz, POz, and Oz. For electrode Pz, we found no main effects (all ps > 0.614) for any of the three factors but a tendency for a two-way interaction of the factors imperative stimulus × door width (F_2,68_ = 2.695, P = 0.075) (no other two two-way interaction effects were significant). However, the interaction of all three factors reached significance (F_2,68_ = 3.199, P = 0.047, partial η^2^ = 0.086). Post hoc comparisons of this three-way interaction showed that in the case of the physical walking condition, there were several significant differences for Go trials for the comparison of Narrow vs. Wide doors (P = 0.004) and Mid vs. Wide doors (P = 0.009), while there were no significant differences between differently wide doors in NoGo trials. In contrast, for the keyboard movement condition, none of the contrasts regarding the different imperative stimuli or door widths reached significance.

For POz, we found a main effect of the factor door width (F_2,68_ = 3.538, P = 0.035, partial η^2^ = 0.094), no impact of the imperative stimulus (F_1,34_ = 1.060, P = 0.310), and no main effect of the movement (F_1,34_ = 0.004, P = 0.950). There was a marginally significant interaction of the factors imperative stimulus and door width (F_2,68_ = 2.794, P = 0.068). The three-way interaction was not significant (F_2,68_ = 2.233, P = 0.115). Post hoc comparisons were done after observing a significant main effect of the factor door width. It showed no significant differences for the comparison of Narrow vs. Mid doors (P = 0.677), but significant differences for the comparison of Narrow vs. Wide doors (P = 0.033), and Mid vs. Wide doors (P = 0.027).

For Oz, we found a main effect of the factor door width (F_2,68_ = 6.744, P = 0.002, partial η^2^ = 0.166), a marginally significant main effect of the factor imperative stimulus (F_1,34_ = 4.019, P = 0.053) but no influence of the movement condition (F_1,34_ = 0.021, P = 0.885). None of the relevant interactions reached significance (imperative stimulus × door width: F_2,68_ = 1.626, P = 0.204; three-way interaction: F_2,68_ = 0.667, P = 0.517). Post hoc comparisons for the different door width conditions showed significant differences for the comparison of Narrow vs. Mid doors (P = 0.010), and Narrow vs. Wide doors (P = 0.002), but not for the comparison of Mid vs. Wide doors (P = 0.337).

#### 3.4.2 PINV

Following EPIC, we noticed a long-lasting negative waveform over central midline electrodes after the onset of the imperative stimulus. As described in previous studies, this waveform represents the postimperative negative variation (PINV), which reflected the immediate motor execution and action readiness related to onset of an imperative stimulus (Casement et al., 2008; Diener, Kuehner, Brusniak, Struve, & Flor, 2009; Elbert, Rockstroh, Lutzenberger, & Birbaumer, 1982; Klein, Rockstroh, Cohen, & Berg, 1996). In the study of Djebbara et al., (2019), the PINV was regarded as reflecting architectural affordances (door width) or, as the authors phrased it, as the sensorimotor dynamics resonating to architectural affordances (Djebbara et al., 2019). To directly compare the PINV observed in the study of Djebbara et al., (2019) with the present study, the PINV component was calculated as the mean amplitude in the range of 600-800 ms after stimulus onset of the imperative stimulus. Both the anterior and posterior PINV components were reported here (Fig. 8).

**Fig. 8.**
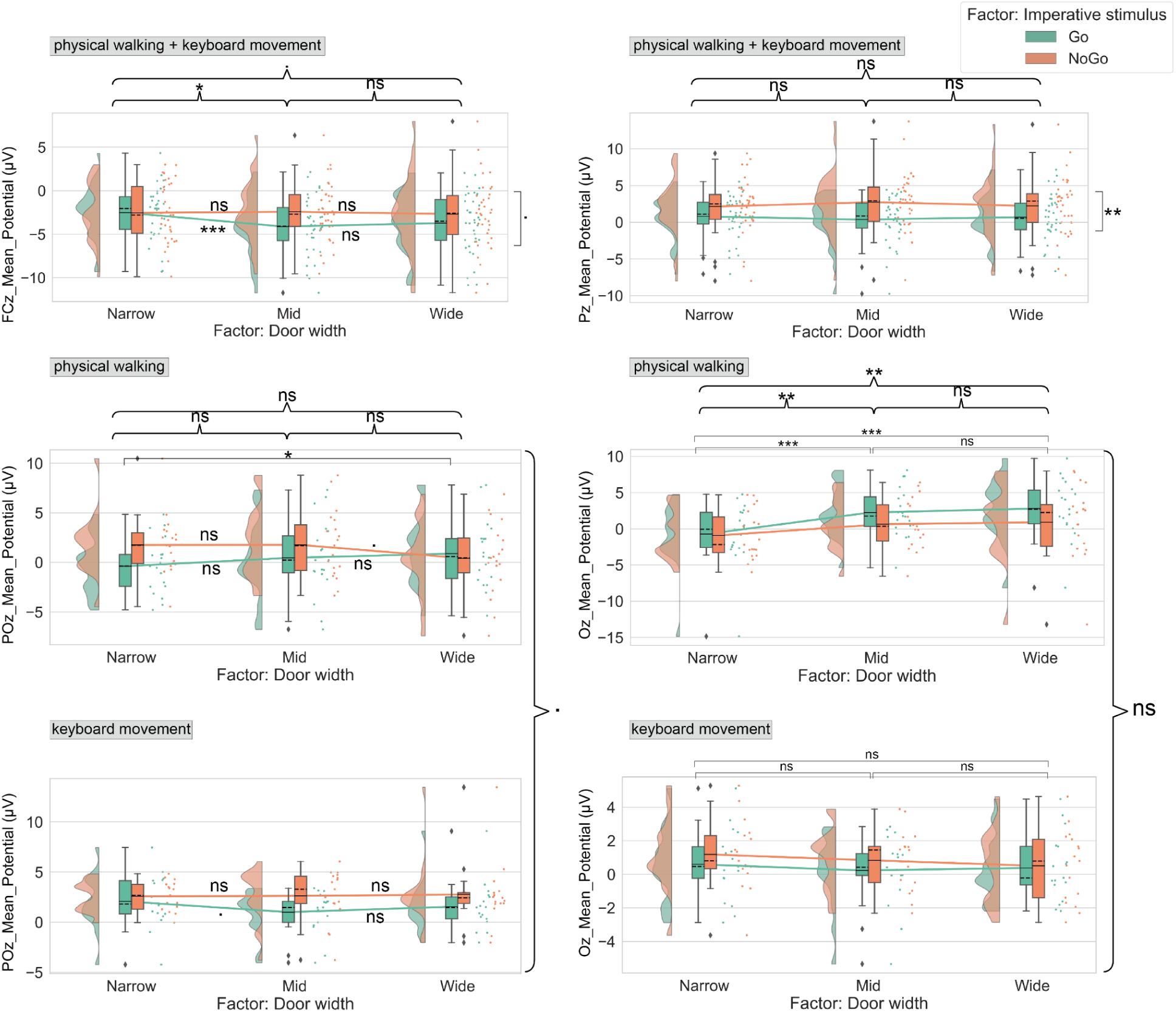
Raincloud plots displaying mean amplitudes of the negative waveform in the time range of 600-800 ms at anterior and posterior electrodes (FCz, Pz, POz, and Oz) for three factors (three different door width conditions (Narrow, Mid, and Wide), imperative stimulus (Go, NoGo) conditions and two movement conditions). The means are indicated by solid lines; medians, by dashed lines. Results of the main effect of the factor door width are indicated by braces, while results of the interaction effect are indicated by square brackets or directly on the solid lines. Results from multiple comparisons using the Tukey’s HSD test and LSD adjustment are indicated with stars. Statistical significance was denoted by ▪P < 0.1, *P < 0.05, **P < 0.01, and ***P < 0.001, while non-significant results were denoted by ’ns’.

##### Anterior PINV - Comparing physical walking and keyboard movement

We conducted a mixed-measures ANOVA with a between-subjects factor movement (physical walking vs. keyboard movement) and two within-subjects factors with 3 (door width) and 2 (imperative stimulus) levels separately for electrodes FCz, Pz, POz, and Oz. For FCz, we found a main effect of the factor door width (F_2,68_ = 3.347, P = 0.041, partial η^2^ = 0.090) and a main effect of the factor movement (F_1,34_ = 9.235, P = 0.005, partial η^2^ = 0.214), and a marginally significant main effect of the factor imperative stimulus (F_1,34_ = 3.666, P = 0.064, partial η^2^ = 0.097). These were qualified by an interaction effect of the factors imperative stimulus × door width (F_2,68_ = 3.563, P = 0.034, partial η^2^ = 0.095). Neither the interaction of the factors door width × movement condition (F_2,68_ = 0.503, P = 0.607) nor the three-way interaction (F_2,68_ = 0.987, P = 0.378) reached significance. Post hoc comparisons of the main effect of the factor door width showed significant differences for the comparison of Narrow vs. Mid doors (P = 0.023), and a tendency of differences for the comparison of Narrow vs. Wide doors (P = 0.071), yet no differences for the comparison of Mid vs. Wide doors (P = 0.775). Post hoc comparison of the two-way interaction effect of the factors imperative stimulus × door width showed that in the Go condition, there were significant differences for the comparison of Narrow vs. Mid doors (P < 0.001) and Narrow vs. Wide doors (P = 0.006); while in the NoGo condition, there were no significant differences between three door width conditions.

##### Posterior PINV - Comparing physical walking and keyboard movement

For Pz, neither the door width (F_2,68_ = 0.064, P = 0.938) nor the movement condition (F_1,34_ = 1.109, P = 0.300) reached significance. However, there was a main effect of the factor imperative stimulus (F_1,34_ = 10.282, P = 0.003, partial η^2^ = 0.232). None of the relevant interactions reached significance (imperative stimulus × door width: F_2,68_ = 1.520, P = 0.226; three-way interaction: F_2,68_ = 0.615, P = 0.544).

For POz, we did not find a main effect of the factor door width on PINV amplitudes (F_2,68_ = 0.043, P = 0.958) but a main effect of the factor imperative stimulus (F_1,34_ = 6.495, P = 0.016, partial η^2^ = 0.160). The factor movement showed a trend toward an effect on PINV amplitudes (F_1,34_ = 3.571, P = 0.067, partial η^2^ = 0.095). Neither the two-way interaction of the factors imperative stimulus × door width (F_2,68_ = 1.333, P = 0.270) nor the two-way interaction of the factors door width × movement conditions (F_2,68_ = 0.903, P = 0.410) reached significance. Finally, a marginally significant three-way interaction was observed (F_2,68_ = 2.629, P = 0.080, partial η^2^ = 0.072). Post hoc comparisons of the three-way interaction showed that in the case of the physical walking condition, significant differences were identified for Go trials for the comparison of Narrow vs. Wide doors (P = 0.036), while there was just a trend of differences for the comparison of Mid vs. Wide doors (P = 0.075) observed in NoGo trials. In contrast, for the keyboard movement condition, none of the contrasts regarding the different imperative stimuli or door widths reached significance, except a trend for the comparison of Narrow vs. Mid doors in Go trials (P = 0.064).

For Oz, we found a main effect of the factor door width (F_2,68_ = 5.993, P = 0.004, partial η^2^ = 0.150), no main effect of the factor imperative stimulus (F_1,34_ = 0.461, P = 0.502), no main effect of the factor movement (F_1,34_ = 0.095, P = 0.760), no interaction effect of the factors imperative stimulus × door width (F_2,68_ = 1.473, P = 0.236), but a significant interaction effect of the factors door width × movement condition (F_2,68_ = 11.669, P < 0.001, partial η^2^ = 0.256). Post hoc comparisons of this two-way interaction effect showed that in the case of the physical walking, there were significant differences for the comparison of Narrow vs. Mid doors (P < 0.001) and Narrow vs. Wide doors (P < 0.001); while in the keyboard movement condition, there were no significant differences between three door width conditions. The three-way interaction was not significant (F_2,68_ = 0.749, P = 0.477).

## 4 DISCUSSION

The main goal of this study was to assess whether and when brain activity underlying affordance perception is altered depending on the kind of interaction with the environment. Such differences would provide further insights into the ongoing debate regarding affordance automaticity. Our assumption was that ERPs under different affordance conditions should vary according to different sensorimotor time windows. Accordingly, we proposed two specific hypotheses: First, early ERPs reflecting different movement conditions should be comparable and therefore independent from the movement context or any other kind of interaction with the environment, reflecting an automated processing of perceived affordance. Second, differences in later motor-related ERPs might be modulated by the movement condition indicating that processing of affordance might be contextually dependent rather than automated at later sensorimotor stages. To examine these hypotheses, we repeated the paradigm of Djebbara et al. (2019) in a condition in which participants moved through the identical virtual space that was used in their study but now using keyboard controls rather than physical walking.

### 4.1 IPQ and SAM data

The analysis of the four dimensions of the IPQ (general presence, spatial presence, involvement, and experienced realism) revealed a significant increase in participants’ sense of presence throughout the experiment, even though their movement through the virtual environment was displayed on a monitor and using keyboard controls. The results indicate that the virtual environment was sufficiently immersive to create a sense of movement through the space allowing for a comparison of the brain dynamics in the present setup with the corresponding data of Djebbara et al. (2019).

The analysis of the SAM ratings for arousal, dominance, and valence consistently revealed significant differences in door width among Go trials, but no differences across NoGo trials for all ratings, except the arousal dimension. Varying door sizes in Go trials consistently yielded differences among all three SAM items, with increasing values observed with increasing door widths. This result is consistent with the finding of Djebbara et al. (2019), except for the dominance dimension. The difference in the latter SAM scale might be the result of additional semantic instructions in the present study to clarify participants’ understanding for the dimension of dominance. In the present study, and in line with the original scales from Bradley & Lang (1994), we used the small SAM manikin representing “being dominated” while the large manikin represented “being dominating”. Overall, these results reflected that our Go/NoGo setting differentially influenced participants’ emotional experience regarding the affordances in the Go and the NoGo conditions. They provided initial evidence from subjective data prior to comparing our motor-related ERPs results with the findings of Djebbara et al., (2019).

### 4.2 Cortical Measures

#### 4.2.1 Early evoked potentials

To examine whether the perception process of affordances is automated during the early sensorimotor time window, we first analyzed the early visual-evoked potentials at posterior leads (Pz, POz, and Oz) comparing the results with the results of Djebbara et al. (2019). We expected to find comparable components and effects of the door widths for ERPs reflecting sensory processing of affordance-related aspects irrespective of the movement condition, i.e., whether participants walked or moved using keyboard controls. Comparable ERP components reflect that affordance perception is independent of movement context, providing evidence of automated affordance perception in early sensorimotor time windows. Overall, the results demonstrated an impact of the door width on amplitudes of the visually evoked components N120, P164, and N260 at posterior leads while no impact of the movement conditions was present (except for the P164 at Pz). Peak amplitudes for the passable doors (mid and wide doors) did not differ significantly, however, peak amplitudes for both door widths were significantly distinct from those for the impassable narrow doors. These findings replicate the results from Djebbara (2019) and align with our first hypothesis regarding the automated perception of affordances during early sensorimotor time windows.

The results demonstrate an association between transition affordances and early visual-evoked potentials. Prior studies on early sensory activity have shown that the visually evoked N1 represents an orienting toward or the engaging of the attentional system (Luck, Heinze, Mangun, & Hillyard, 1990), especially in the case of involuntary visual attention (Escera, Alho, Winkler, & Näätänen, 1998; Escera, Yago, & Alho, 2001; Näätänen, 1990; Wang, Wu, Fu, & Luo, 2010). The P1, on the other hand, represents a facilitation of early sensory processing for stimuli occurring in the attended location (Luck et al., 1990). The N2-component represents attentional selection of relevant information, voluntary attention, and the capability of inhibiting distraction (Luck & Hillyard, 1994; Patel & Azzam, 2005; Schneider, Beste, & Wascher, 2012; Y. Wang, Wu, Fu, & Luo, 2010), as well as motor anticipation or prospective movement (Kourtis, Sebanz, & Knoblich, 2012). The amplitudes of the early visual evoked N1-P1-N2 found in the current study thus might reflect perception of architectural affordances as a transition from a stimulus-driven process to a goal-directed perceptual process involving voluntary attention and motor anticipation.

The direct statistical comparison of the current posterior N1-P1-N2 components with the corresponding results of Djebbara et al. (2019) revealed no major differences between both movement conditions in the early sensorimotor time window with the exception of a main effect of the movement context at Pz for the P164. The P164 did not vary depending on the door widths nor did the interaction of the two factors door width and movement conditions reach significance. Therefore, the stronger peak amplitudes of P164 at Pz observed in the physical walking condition seem to reflect a generally enhanced supramodal activity, potentially resulting from increased immersive optic flow in a head-mounted VR environment (Li et al., 2020). The absence of a significant interaction effect movement context × door width supports the assumption that early perception stages of affordance processing are not influenced by the movement context.

Overall, these findings show that the processing of architectural affordances is associated with differences in attentional processes during early sensorimotor time windows but that these are independent from how participants move through virtual space. Our results suggested that the early perceptual processes regarding affordances are independent from the movement context and, thus, reflect an automated process.

#### 4.2.2 Motor-related potentials

To further investigate whether a later sensorimotor time window also reflects automated affordance processes or rather context-dependent modulations, we analyzed motor-related potentials in the keyboard-controlled movement condition and compared them with the results of Djebbara et al., (2019).

Regarding the factor door width, we found significant differences in amplitudes of the posterior EPIC at Oz and POz and also the anterior EPIC at FCz; yet for these electrodes and also for electrode Pz, there were no significant interactions of imperative stimulus × door width demonstrating consistent results in both studies. The nature of the EPIC component has not been as well investigated as other ERP components. Therefore we attempted to discuss the role of EPIC by combining our experimental paradigm and EPIC statistical data. In our experiment, after viewing the display of the transition for 6 seconds (σ = 1), participants would see the color of the transition door turned from gray to either green or red. According to this imperative stimulus, they decided whether to pass through the door or turn their body to rate the SAM questionnaire. And EPIC was time-locked with the onset of the imperative stimulus. As we mentioned above, in the present desktop movement condition, we found significant impacts of different door widths on the EPIC amplitudes at electrodes FCz, POz and Oz. It might reflect participants’ increasing attention in differentiating imperative signals while considering varying affordances. In view of the latency differences between the EPIC and PINV components in our study, the imperative differentiation of EPIC should occur at an intermediate stage of affordance perception after the early visual-attention and before subsequent motor execution processes.

Following the EPIC, a prominent PINV component was observed, indexing motor execution and control processes (Djebbara et al., 2019; Elbert, Rockstroh, Lutzenberger, & Birbaumer, 1982). At the anterior electrode FCz, consistent with the findings of Djebbara et al. (2019), a two-way interaction effect of the factors door width × imperative stimulus was observed, with no interaction regarding the factor movement. However, the impact of the movement context became evident at posterior leads covering cortical regions related to the visual processing. Here we found interaction effects related to the movement context. The context-dependent modulation of components reflecting perceptual affordance processing during late sensorimotor time windows support our second hypothesis. Specifically, the two-way interaction of factors door width × movement condition reflected that during later sensorimotor time windows, motor-related brain dynamics (PINV) regarding varying transition affordances were modulated by movement context. Variations in PINV amplitudes for varying affordances were amplified under the physical walking condition. Furthermore, the trend toward a three-way interaction showed that this amplification should be attributed to Go trials, rather than the NoGo trials. Generally, in Go trials participants were instructed to pass the door and the physical walking condition was the more natural condition and thus arguably more beneficial for this kind of agent-object interaction. Therefore, for the PINV, an interaction of the door widths with the movement condition might be attributed to the limitations of the current keyboard movement condition reflecting a less embodied way of interacting with the environment. The trend of the PINV reflected that participants might have a decreased sense of control underlying their anticipated motor trajectory towards unpassible transitions (Djebbara et al., 2019). However, the post hoc comparisons of the interaction effect in the keyboard setup revealed no significant differences for differently wide doors, even though the ERP plots indicated an affordance trend of the PINV.

Regarding the above post hoc comparisons of the three-way interaction, we considered that in “Go” trials, perception was shaped by the anticipated motor trajectory, which varies based on the transition affordances. However, this shaping was also constrained by the movement condition, that is, either physical walking through the VR environment or using keyboard controls. In the current keyboard movement condition, participants’ bodily movement was not allowed, therefore their embodied information and proprioceptive feedback might also be limited. Therefore, in the Go condition, although they were shown failed feedback when they passed the unpassible doors, their motor anticipation of the trajectory to transit might not be strong enough to elicit significant differences of cortical potentials among three door widths. Accordingly, although the PINV findings of Djebbara and colleagues (2019) suggested there were significantly different senses of loss of control between three door widths in the Go trials, we can not get this inference for certain in the current case (keyboard movement). However, in “NoGo” trials, there were no significant impacts of door width on the PINV component, from both the results of Djebbara et al., (2019) and the current datasets. This reflected that in this case, perceptual processes might not depend on the movement context. Different from the psychological processes related to motor execution and motor control under the Go condition, in cases of NoGo, participants did not need to interact with the transition, but rather simply turned back to answer the questionnaire. In other words, the brain dynamics data under the NoGo condition might not be attributed to the anticipated motor trajectory based on proprioceptive information of varying transition affordances, but rather homogeneous, simple and immediate body-turning execution, which was independent of the movement.

Besides the above context-dependence modulation findings of affordance perceptual processing, we also found an interaction of door width and imperative stimuli at the electrode FCz, which was consistent with the results of Djebbara et al., (2019). This might indicate that motor-related brain dynamics regarding varying affordances at the motor-related cortical regions tends to keep consistent between different movement contexts, even during the late sensorimotor time window. This suggests that besides the consideration of the time window, there exist more potential modulators regarding varying affordance contextual dependence, such as cortical regions, etc.

Taken together, only when participants knew that they would interact with doors of different transition affordances (Go trials), sensorimotor brain dynamics time-locked with the onset of the imperative stimulus covaried with the door widths reflecting whether the door was perceived as passable or not. Additionally, this covariation of the PINV with differently wide doors was found only when participants were physically walking through the space, consistent with our second hypothesis. This supports the assumption that active exploration influences the emergence of affordances, which in turn shapes and affects the agents’ perception and exploration of an environment (Wang, Sanches de Oliveira, Djebbara, & Gramann, 2022). Last but not least, our findings regarding context-dependent modulations of the EPIC and PINV components might reflect multiple subphases of affordance perception, including both stimulus-driven and goal-directed processes.

## 5 Conclusion

In comparison to the physical walking condition in the study by Djebbara and colleagues (2019), the current study used keyboard controls to move through the identical virtual environment. The two experiments revealed a significant impact of the door width on early sensorimotor processes as reflected in amplitude modulations of early ERPs, while later motor-related components were dependent on the movement context. The evidence from electroencephalography (N1-P1-N2 amplitude) showed two types of processes, namely, involuntary attention and voluntary attention in sequence, which might reflect the transition from an stimulus-driven process (bottom-up) to a goal-directed process involving anticipation (top-down). In the late sensorimotor time window, the brain dynamics data showed two sets of main psychological activities: imperative differentiation, and subsequent motor anticipation, execution, and control. As affordance processing progressed, it gradually involved more sub-processes that required greater top-down control. This progression reflected the sensorimotor integration underlying processing of perceived affordance and a strong dependence on the context, which is in line with the account of the SMCs scheme (Djebbara et al., 2022) and Hierarchical Affordance Competition (Cisek, 2007; Pezzulo & Cisek, 2016). However, we note that early and late stages of affordance processing should not be seen as separate processes but rather as points on a continuum, with early affordance perception being the first step in a sequence of stages leading to late affordance perception-action coupling. Notably, the transition from the early, stimulus-driven process to the late, goal-directed, sensorimotor integration processes requiring greater top-down control may occur gradually, even as early as the late latency of the early visual attention stage and the subsequent imperative differentiation stage.

## AUTHOR CONTRIBUTIONS

S.W., Z.D., and K.G. designed research; S.W. and K.G. developed the research question, hypotheses and conceptualization; S.W. performed research; S.W. and K.G. analyzed data; and S.W., Z.D., G.S.O. and K.G. wrote the paper (S.W. and K.G.: Original draft; Z.D., G.S.O. and K.G.: Review & editing).

## ACKNOWLEDGMENTS

The authors would like to thank our engineer Benjamin Paulisch for his Unity programming, our student assistants Yiru Chen for her support in conducting the experiment and S.W.’s academic peer Zhenxing Hong (National Cheng Kung University, Tainan, Taiwan) for his assistance in enhancing the graphic aesthetics of the present study. Z.X.H. adeptly combined sub-plots, added informative arrows and statistics, and skillfully improved the figures using Adobe Illustrator.

