## Supplemental figure S1. S2. S3 for "Human Brain Dynamics Dissociate Early Perceptual and Late Motor-Related Stages of Affordance Processing"

* Sheng Wang

Zakaria Djebbara

Guilherme Sanches de Oliveira

Klaus Gramann

This PDF file includes:

Fig. S1: ERP plots for MRCPs at 6 electrodes (Fz, FCz, Cz, Pz, POz, Oz)

Fig. S2: Rain-cloud plots for component EPIC at 6 electrodes (Fz, FCz, Cz, Pz, POz, Oz)

Fig. S3: Rain-cloud plots for component PINV at 6 electrodes (Fz, FCz, Cz, Pz, POz, Oz)

### Supplementary Information Text

Subhead. Additional event-related potential plots of all 6 electroencephalograph channels for MRCPs. Rain-cloud plots of contrasts for the components EPIC and PINV included.

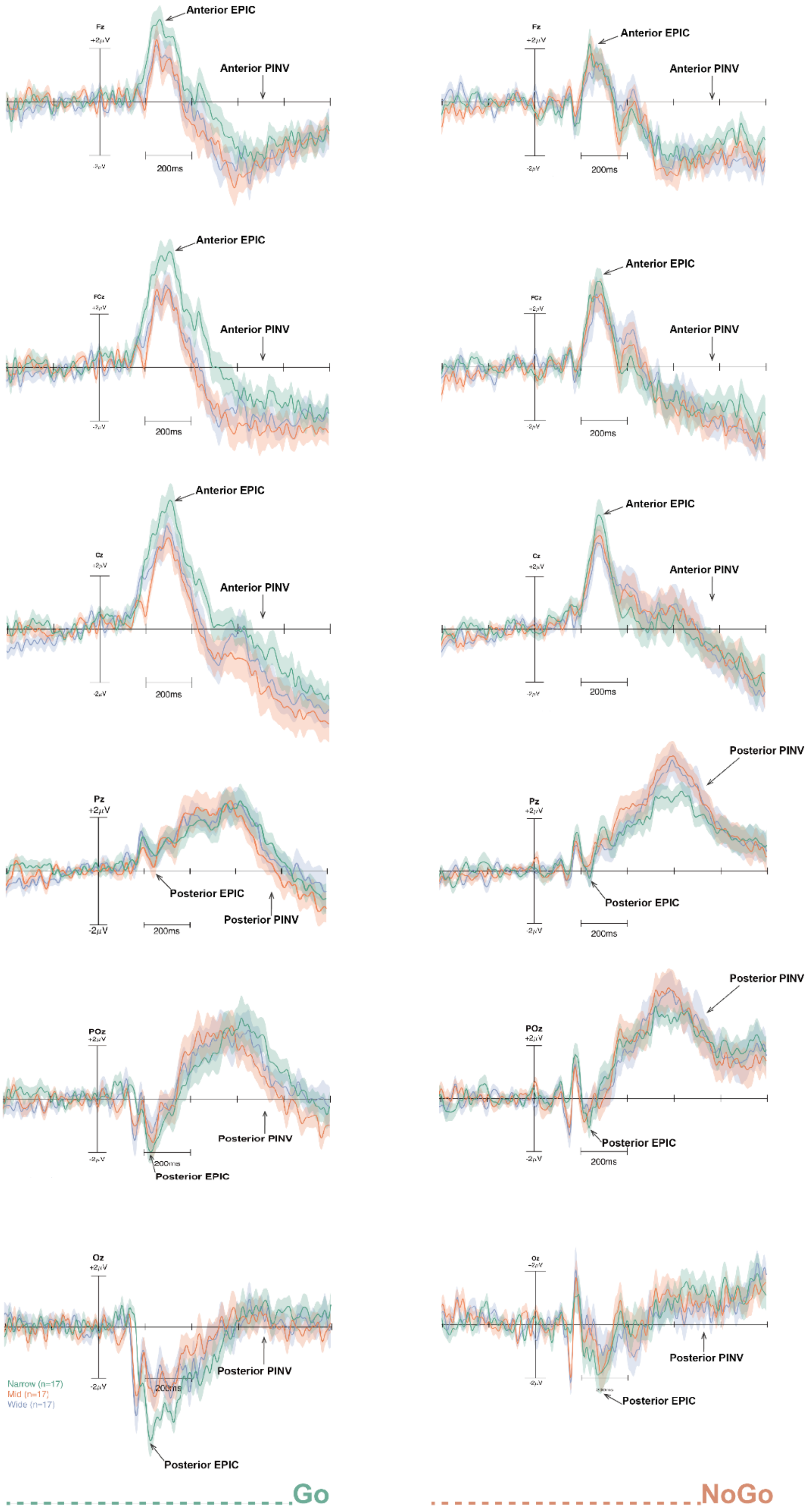

Fz
+2μV
-2μV
200ms
Anterior EPIC
Anterior PINV
FCz
Cz
Pz
POz
Oz
Posterior EPIC
Posterior PINV
Narrow (n=17)
Mid (n=17)
Wide (n=17)
Go
NoGo

**Fig. S1.** Six time-locked event-related potentials (ERPs) from electrode Fz, FCz, Cz, Pz, POz, and Oz at the onset of imperative stimulus (Go, NoGo) under the present keyboard movement condition. The Narrow condition is in green, the Mid condition is in red, and the Wide condition is in purple. The plots on the left column showed the ERPs under the Go condition, while the plots on the right column showed the ERPs under the NoGo condition. Two components (EPIC and PINV) were marked with arrows.

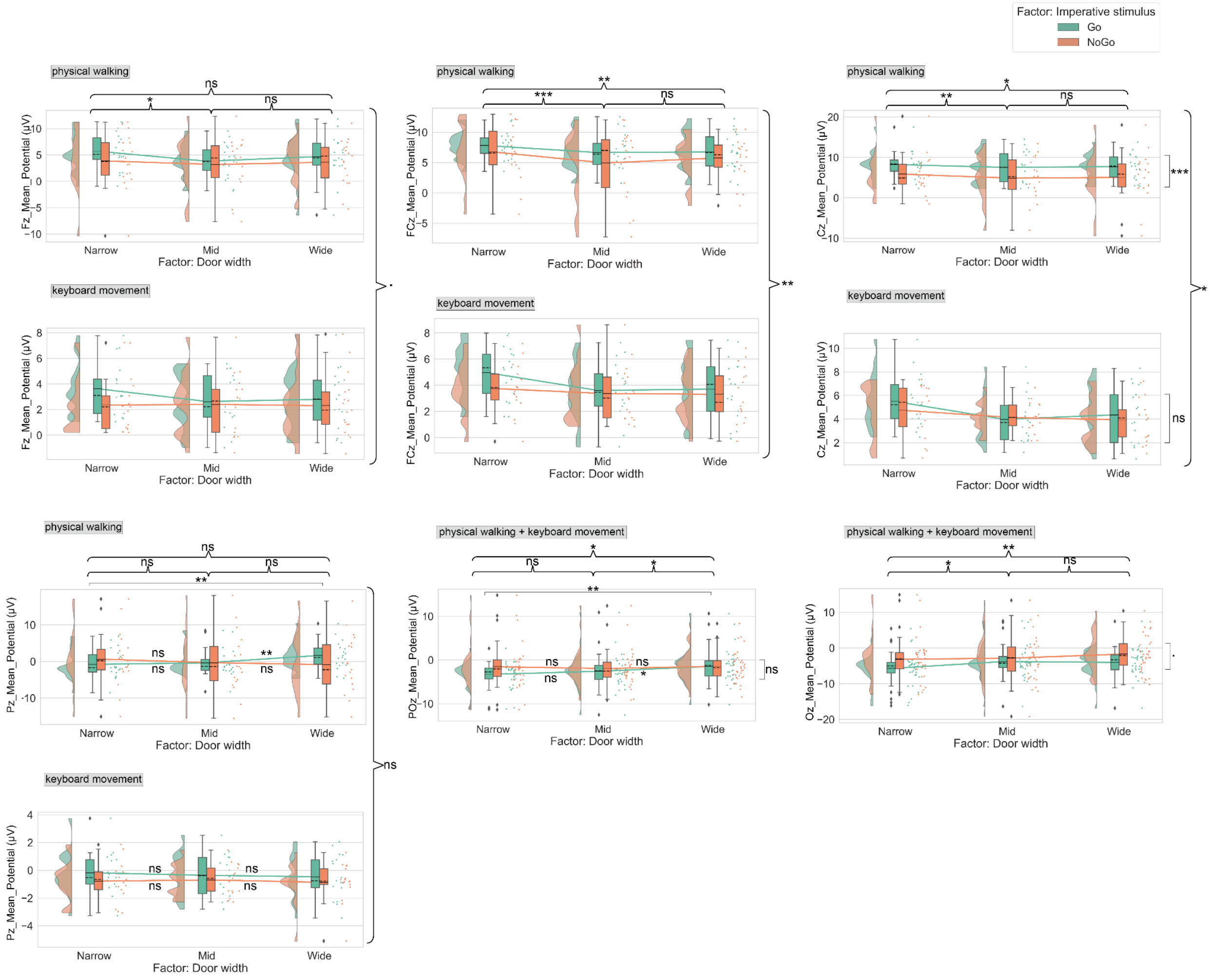


**Fig. S2.** Raincloud plots displaying mean amplitudes of EPIC peaks at anterior and posterior electrodes (Fz, FCz, Cz, Pz, POz, and Oz) for three factors: three different door width conditions (Narrow, Mid, and Wide), imperative stimulus conditions (Go, NoGo), and two movement conditions (physical walking and keyboard movement). The means are indicated by solid lines, and medians by dashed lines. Results of the main effect of the factor door width are indicated by braces, while results of the interaction effect are indicated by square brackets or directly on the solid lines. Results from multiple comparisons using the Tukey's HSD test and LSD adjustment are indicated with stars. Statistical significance was denoted by ▪$P < 0.1$, *$P < 0.05$, **$P < 0.01$, and ***$P < 0.001$, while non-significant results were denoted by 'ns'. Generally, for the additional electrode Fz, we observed a tendency for a main effect of the factor movement ($F_{1,34} = 3.843$, $P = 0.058$). For the additional electrode Cz, we observed a main effect of the factor door width ($F_{2,68}$ = 5.528, P = 0.006, partial $\eta^2$ = 0.140), a main effect of the factor movement ($F_{1,34}$ = 4.373, P = 0.044, partial $\eta^2$ = 0.114) and an interaction of factors imperative stimuli × movement conditions ($F_{1,34}$ = 6.176, P = 0.018, partial $\eta^2$ = 0.154).

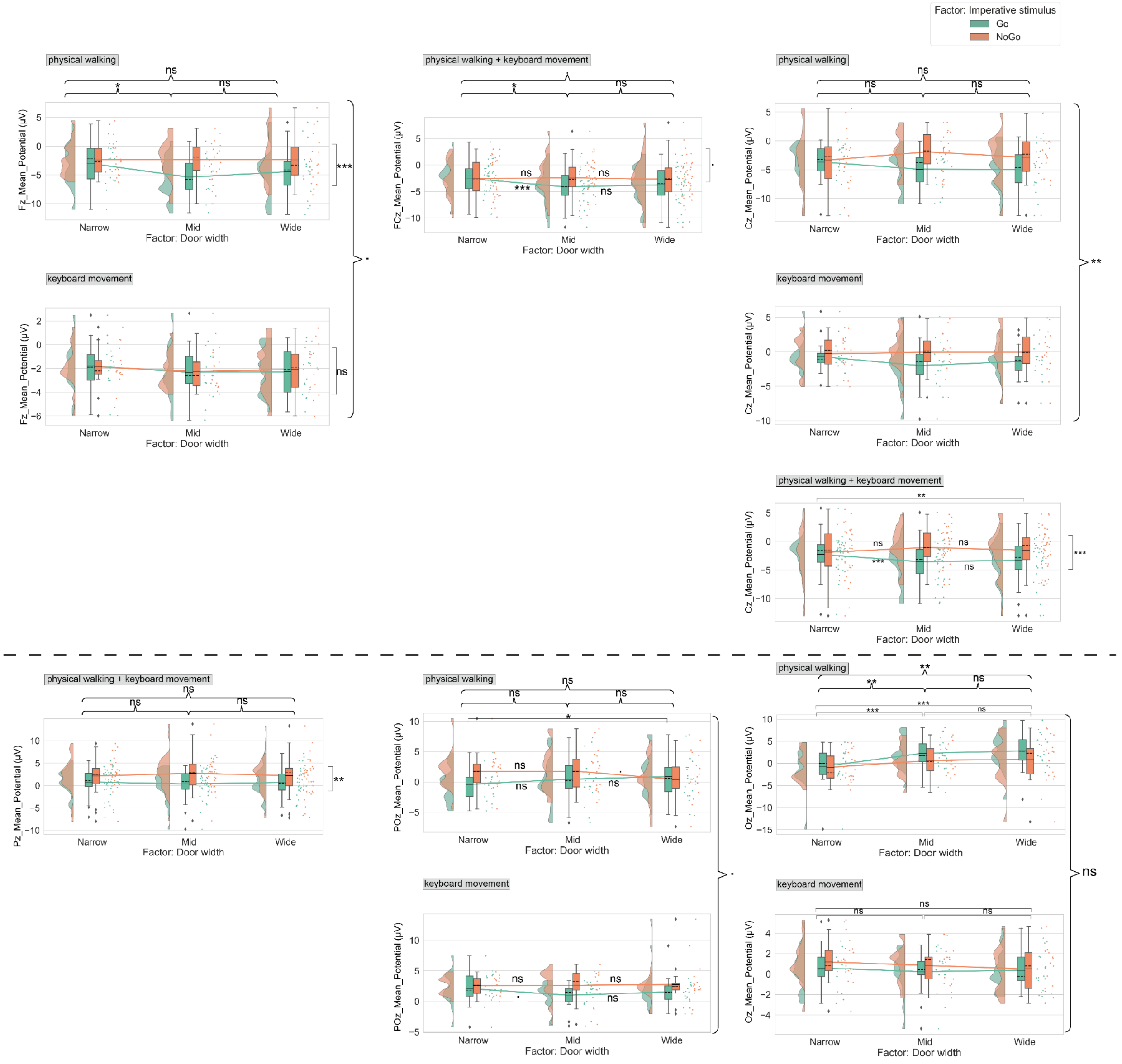

**Fig. S3.** Raincloud plots displaying mean amplitudes of the negative waveform in the time range of 600-800 ms at anterior and posterior electrodes (Fz, FCz, Cz, Pz, POz, and Oz) for three factors: three different door width conditions (Narrow, Mid, and Wide), imperative stimulus conditions (Go, NoGo), and two movement conditions (physical walking and keyboard movement). The means are indicated by solid lines, and medians by dashed lines. Results of the main effect of the factor door width are indicated by braces, while results of the interaction effect are indicated by square brackets or directly on the solid lines. Results from multiple comparisons using the Tukey's HSD test and LSD adjustment are indicated with stars. Statistical significance was denoted by ▪P < 0.1, *P < 0.05, **P < 0.01, and ***P < 0.001, while non-significant results were denoted by 'ns'. Generally, for the additional electrode Fz, we observed an interaction of factors imperative stimuli × movement conditions ($F_{1,34} = 6.481$, $P = 0.016$, partial $\eta^2 = 0.160$). For the additional electrode Cz, we observed a main effect of the factor movement ($F_{1,34} = 8.913$, $P = 0.005$, partial $\eta^2 = 0.208$), and an interaction of factors imperative stimuli × door width ($F_{2,68} = 5.308$, $P = 0.007$, partial $\eta^2 = 0.135$).
